# Toward an Objective Assessment of Fight-or-Flight System Function in Anxiety Disorders: Startle Reflex Modulation in Threat Coping Contexts as a Candidate Neurophysiological Marker of Disrupted Defense System Operation

**DOI:** 10.64898/2026.09.16.751525

**Authors:** Joshua H. Gertler, Christina Marsicano, Mark S. George, Lisa M. McTeague, Christopher T. Sege

## Abstract

**Background:** As a key mechanism of disorder, clinical anxiety changes how a defensive motivation system functions in active threat coping contexts. Building on initial evidence that a lab-based escape/avoid preparation task is sensitive to disruptions in underlying defensive activation, current research moved toward implementation as a clinical assessment by testing if task responding predicts symptoms in anxiety treatment seekers.

**Methods:** In our escape/avoid task, unpleasant images are shown after 5.5-second cues that indicate if the upcoming image can be prevented (“avoid” context), terminated after onset (“escape” context), or not controlled (“no-control” comparator). To assess defensive activation during response preparation/threat anticipation, a startle reflex indicator of such activation is probed during each cue. Guided by prior data, 125 individuals were classified into groups based on whether startle was enhanced 1) in a no-control, relative to avoid, context, and 2) in an escape, relative to avoid, context. To test clinical correlates, anxiety (State-Trait Anxiety Inventory), depression (Beck Depression Inventory), and worry (Penn State Worry Questionnaire) were then compared across response groups.

**Results:** Anxiety and worry were higher in participants with enhanced no-control *and* escape startle than in participants with enhanced no-control but *not* escape startle. Further, depression was higher in participants with *under-enhancement of no-control startle* than other groups. Finally, when comparing startle-based profiles to canonical questionnaire cut-off score-based groups, ∼54-63% convergence emerged.

**Conclusions:** Defensive profiling in escape/avoid contexts shows promise as a prognostic indicator, but a need to continue improving is also indicated. Possibilities to improve predictiveness and reliability are discussed.

## Introduction

In adults who receive frontline cognitive-behavioral treatments in the United States, remission rate for anxiety disorders overall is estimated at 51%,^1^ and for related post-traumatic stress or obsessive-compulsive disorders at 53%^2^ and 40%^3^ respectively. Given variable outcomes, much research has tested readily observed baseline variables (e.g., demographic and self-reported clinical variables) as treatment response predictors and found some prognostic value.^4,5^ At the same time, there is much room for improvement – and because self-report variables in particular have validity-limiting features (e.g., recall/ reporting bias, indirectness as indices of underlying pathologic processes^6^), one proposed approach to improving prognostic capability is by identifying direct neurobiological (*objective*) indicators of underlying psychiatric disorder-driving mechanisms. This approach is central to Research Domain Criteria and other initiatives that seek to create a precision medicine of psychiatric practice^7,8^ – bringing psychiatry closer to other fields of medicine that use validated biological markers of underlying pathogens to guide treatment that then directly targets those causal pathogens rather than overlying symptoms.

In anxiety, traumatic stress and obsessive-compulsive disorders, disrupted operation of a defense (“fight-flight-freeze”) system in threat contexts is a particularly crucial disorder-driving process and, thus, treatment target.^9,10^ Defense system dysfunction is especially critical inasmuch as it motivates exaggerated reliance on escape/ avoidance behavior as a coping strategy – which then becomes a key driver of impairment. Toward an ability to objectively assess underlying defensive dysregulation as an escape/ avoidance motivator, Sege and colleagues^11-13^ have built on earlier avoidance response preparation designs^14-16^ to develop a task wherein defense system operation is objectively measured across contexts with varying levels of active control over upcoming threat. Briefly, this task presents unpleasant images under conditions in which: a) a quick button press *before image onset* prevents the image (“avoid” context), b) a quick button press *after image onset* removes the image (“escape” context), or c) there is no control over how long the unpleasant image is presented for (“no control” context). By measuring defensive activation during pre-picture cues that indicate the level of control over upcoming exposure, this task assesses integrity of defensive *regulation across threat coping contexts* – guided by contemporary preclinical and translational models that describe how subtle disruptions in defensive regulation across contexts are likely more critical to anxiety and related disorder etiology than are gross, context-insensitive defensive response exaggerations.^17-19^

To date, defensive activation in the escape/ avoidance preparation task has been assayed with an established neurobiological index of such activation, the blink startle reflex.^20^ Startle reflex methodology involves presenting brief, rapid-onset (typically auditory) stimuli to prompt a series of motor reflexes during a task and then measuring how a component of that series (often the reflexive eyeblink) is modulated by foreground task demands.^20^ Measuring startle during emotional processing tasks has consistently demonstrated that the startle blink is modulated by emotional state^21^ – including with reliable *priming* of the blink reflex (a defensive reaction that protects the eye from sudden damage^20^) observed during unpleasant image viewing.^22^ Concurrently, circuit delineation has shown that blink startle reflex modulation is produced by circuit interactions between the amygdala and brainstem (i.e., periaqueductal grey),^23^ which directly modulate a simple startle reflex circuit by inputting to its nucleus reticularis pontis caudalis node.^24,25^ Supported by these data, then, startle blink reflex modulation can be measured as a direct probe of defensive system operation during threat processing tasks.

With startle as an index of defense system regulation in the escape/ avoidance task, our studies to date then reveal typical reflex priming (*potentiation*^20^) during no-control threat anticipation *and also some potentiation during escape, as compared to avoid, preparation* at the group level.^11,12^ Additionally, a study examining individual differences in an analogue sample found a relationship between trait-level anxiety and startle potentiation *in escape (but not avoid or uncontrollable) contexts*.^12^ Following this up, a subsequent study with a new non-treatment-seeking sample (n=20) and a sample of clinically anxious treatment seekers (n=25) found that non-treatment-seeking subjects showed similar startle *inhibition* (an indicator of instrumental task engagement^26^) in avoidance and escape contexts (along with reflex potentiation in a no-control context), but treatment seeking participants, instead showed reflex inhibition only in the avoid context and potentiation in no-control *and escape* contexts – suggesting a *context-specific* insensitivity to *escape* response availability in clinically anxious individuals.^13^ Further extending analogue sample findings, regression analyses in the clinical sample also revealed that, while trait anxiety correlated with increased startle in an escape context, *depressive* symptomatology (as measured by the Beck Depression Inventory, BDI^27^) correlated with *reduced reactivity <u>in a no-control condition</u>* – an effect apparently driven by 5 individuals who reported severe (BDI>29^28^) depression and did not show modulation in any context. Taken as a whole, then, findings to date provide initial evidence for reliable sensitivity of an escape/avoid task to 1) general defense system regulation across active threat coping contexts, and 2) distinct patterns of clinical disruption in this regulation in anxious versus severely depressed individuals.

To build on initial evidence of escape/avoid task sensitivity to clinically relevant changes in defense system function, one critical next step will be to replicate results in larger clinical samples – especially given the small severely depressed sample in preliminary work. In addition, with a long-term objective to replace subjective data with biological measures as prognostic indices, testing the degree to which clinical presentation can be determined by classifying individuals based on their objective data – rather than the other way around – is another key research step.^29^ With this in mind, the current study expanded our sample of mental health treatment seeking individuals to: 1) examine if profiling individuals based on reflex priming under threat (i.e., escape and no-control) conditions reveals measurable symptom differences, and; 2) test correspondence of startle reflex profiles with traditional symptom-based profiles. Based on prior data, we predicted that: 1) classification of escape potentiation would reveal groups with similar startle in avoid and no-control contexts but who differ specifically in whether reflexes in an escape context are inhibited (“action-sensitive” group) or primed (“threat-sensitive” group), and; 2) classification of no-control potentiation would reveal a third (“context-insensitive”) group with overall minimal startle modulation. With these groupings, we then predicted that a threat-sensitive group would report higher anxiety than an action-sensitive group, and a context-insensitive group would report higher depression than other groups. Further, we predicted correspondence between startle-based and symptom-based profiling approaches – such that, when individuals are classified by canonical clinical cut-off scores as in prior work, those with clinically heightened anxiety would show a threat-sensitive startle modulation profile and those who also have severe depression would show a context-insensitive profile.

## Methods and Materials

### Participants

Participants were recruited from outpatient mental health clinics in a Southeastern United States medical center and the surrounding community. Participants were recruited after initially presenting to clinic for treatment for an anxiety or related disorder, or by flyers in the area surrounding the medical center. Study procedures were approved by the institutional review board, and all participants provided written consent. To maximize power for this new analysis, the current sample combines data from 90 newly recruited anxiety treatment-seeking adults with data from the 25 anxiety treatment-seeking adults and 20 community-dwelling non-treatment-seeking (at time of participation) adults who participated in our prior work.^1^ All participants in this research were between the ages of 18–65.

### Study Task

All participants completed an experimental task developed in our prior work that presents visual cues and then highly unpleasant pictures as response-motivating outcomes (**Figure 1**; see Sege et al., 2018^11^ for further task details). On each trial, an initial 5-to 5.5-second predictive cue is immediately followed by a 500ms visual go signal (on avoidance trials) or unpleasant (disgusting/ violent) picture (on escape and no-control trials). On avoid and escape trials, if participants press a button within 500ms after cue offset, the go signal or aversive picture is replaced by a neutral (everyday scene/object) image that is presented for 2.5 more seconds. Conversely, if a button is not pressed within 500ms on avoidance trials the go signal is replaced by an unpleasant picture that is then shown for 2.5 seconds, or on escape trials unpleasant exposure continues for another 2.5 seconds. On no-control trials, initial 500-ms unpleasant exposure is followed by 2.5 seconds more exposure regardless of any button presses (no-control trials with button press removed; <1% of all trials).

**Figure 1.**
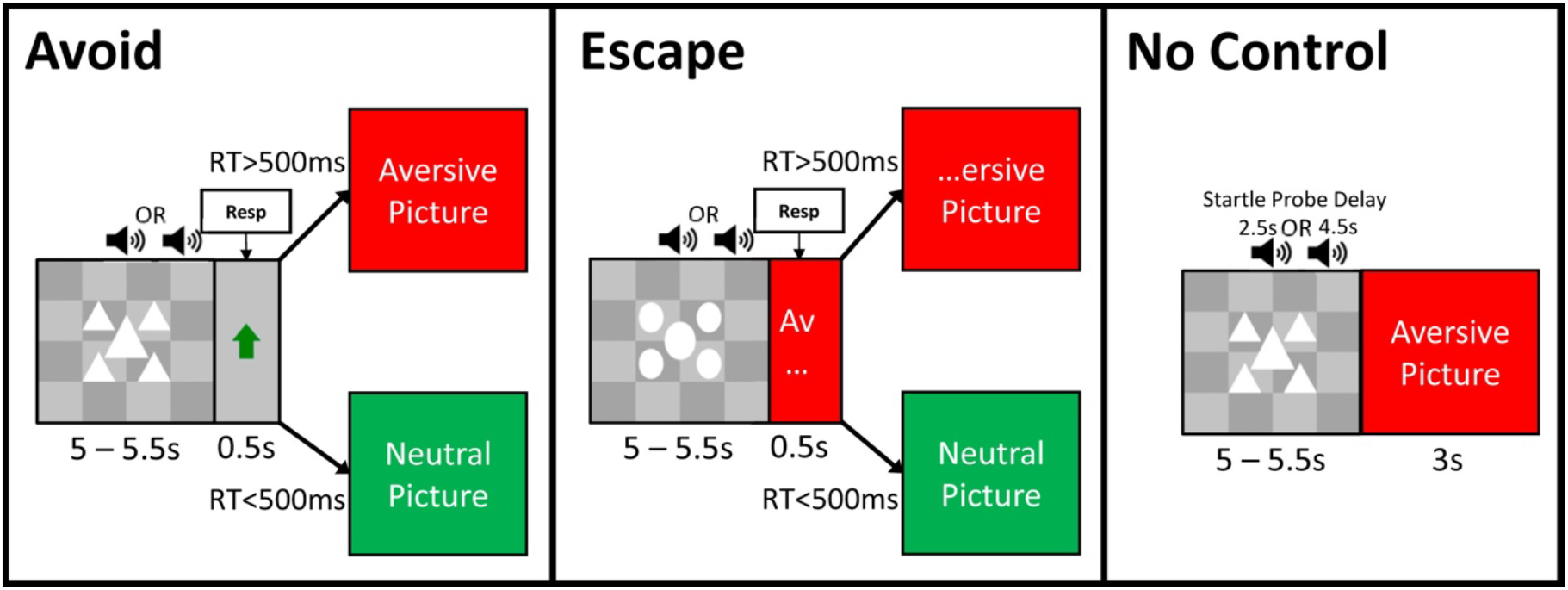
Task design including actual cues, go signal, and timings. Resp = response; RT = response time.

Task orders were counterbalanced across participants such that each image occurred in each context (no image repetition). Avoid, escape, and no-control trials are presented in pseudorandomized event-related (i.e., not blocked) order such that each trial type could occur no more than twice consecutively. In addition to visual stimuli, a 50-ms, 105-dB white noise burst was presented during each cue (2.5 seconds or 1 second before anticipated stimulus onset) to probe startle reflex reactivity.

### Sample Characterization Measures

Participants also completed clinical characterization questionnaires which included: 1) a demographics survey; 2) the State-Trait Anxiety Inventory – Trait Anxiety index (STAI-T);^30^ 3) the BDI – Second Edition (BDI-II);^27^ 4) an abbreviated Penn State Worry Questionnaire (PSWQ-A);^31^ 5) a short-form Intolerance of Uncertainty Scale (IUS-12);^32^ 6) the Anxiety Sensitivity Index-3 (ASI-3);^33^ 7) the PTSD Checklist for DSM-5 (PCL-5);^34^ 8) a Brief Emotional Avoidance Questionnaire (BEAQ),^35^ and; 9) an Illness Intrusiveness Rating Scale(IIRS)^36^ as a survey of mental health–related functional interference. In addition, psychiatric diagnosis was assessed by doctoral-level staff via a Mini-International Neuropsychiatric Interview.^37^

### Procedure

After interviews and surveys, participants were seated in front of the task computer and affixed with sensors, headphones, and a mouse to respond. Before the task, subjects were told each cue’s meaning and told: 1) not to press the button on uncontrollable trials; 2) that they would lose control over the unpleasant image if they pressed the button *before* cue offset (to discourage early and/ or repeated responding); and 3) to ignore noises played over headphones. Participants then completed 5 practice trials to verify understanding and present 5 startle habituation probes.^38^

After task completion but before sensor removal, participants rated the pleasantness of anticipating avoidable, escapable, and uncontrollable aversion and of viewing disgusting/violent and everyday images using a Likert-type scale ranging from 1 (very unpleasant) to 9 (very pleasant; 5 indicates neither pleasant nor unpleasant). Sensors were then removed, and subjects were debriefed and paid.

### Blink Startle Reflex Measurement and Processing

Startle blink reactivity was measured using an MP150 data acquisition system and 2 Ag/ AgCl sensors measuring electromyographic signal in the orbicularis oculi (blink) muscle under the left eye. Sensors were connected to an EMG100C amplifier (BIOPAC Systems, Inc), filtered online at 10-500Hz, and sampled at 1000hz. Offline, after rectification and low-pass 40hz filtering,^38^ blink responses to each probe were scored and standardized as *T* scores for each subject (relative to mean/ standard deviation of all blinks for that subject).^38^

Guided by previously observed patterns,^11-13^ after data collection subjects were classified based on Escape – Avoid and No Control – Avoid threat potentiation scores. As a first step, subjects were grouped based on whether their No-Control – Avoid score was greater than 1 *T* score point – indicating normative potentiation during uncontrollable threat anticipation. As a second step, participants with normative no-control potentiation were then classified based on whether their Escape – Avoid was also greater than 1 *T* score point.^2^ This resulted in three profiles of reflex modulation – a profile of typical potentiation in a no-control context but not in an escape context (“action-sensitive”), a profile of potentiation in no-control *and* escape contexts (“threat-sensitive”), and a profile where participants atypically did not have startle potentiation during unpleasant anticipation (“context-insensitive”).

### Data Analysis

Data were prepared for analysis in JmP 19 software©, and analyses were conducted using JASP© software. For questionnaires, missing items were imputed with JmP’s Automated Imputation function that imputes missing values via a low-rank matrix approximation method that automatically selects the best dimension for imputation based on available data for the survey that includes the imputed item.

Following data processing/ scoring, main analyses involved analyses of variance (ANOVAs) with Startle Group (action-sensitive, threat-sensitive, context-insensitive) as a between-subjects factor and each questionnaire score as an outcome in separate analyses for each survey. In these analyses, a significant effect of Startle group was followed up with Bonferroni-corrected (n tests=3) independent-samples *t*-tests. In addition, differences in the pattern of startle modulation for each startle group were examined using Bonferroni-corrected paired-samples *t*-test context comparisons within each group (n tests=3) and independent-samples *t*-test comparisons of each context across groups (n tests=3). Finally, groups were also compared for age using the same approach as for questionnaire scores, and for biological sex at birth and dichotomized (minoritized vs. non-minoritized) racial/ ethnic identification via Fisher’s Exact *χ*^*2*^ tests.

Complementing tests of clinical differences as a function of startle reflex profiling, we also replicated our prior work that classified participants based on suggested clinical cut-oas for elevated trait anxiety (STAI-T score > 39^11,39^) and severe depression (BDI-II score > 29^28^). Because of observed differences in worry across startle groups (see **Results**), clinic-level worry (PSWQ-A ≥ 22^40^) was also explored as an additional criterion. Following symptom-based classification, differences in age, biological sex at birth and dichotomized race/ ethnicity identification were done as for Startle Group analyses, and mixed-eaects ANOVA with Clinical Group (Subthreshold, Worry Only, Clinical Anxiety, Severe Depression; see **Results**) as a between-subjects factor and Task Context (Avoid, Escape, No-Control) as a within-subjects factor^41^ were then done to test for differences in startle modulation across clinical groups. Follow-up of a significant Group X Context interaction was with Bonferroni-corrected (n tests=3) paired-samples *t*-test comparisons of contexts within each group, and (n tests=4) independent-samples *t*-test comparisons of groups within each context.

For each grouping described above, any group differences in age, sex at birth or race/ ethnicity identification led to that variable’s inclusion as a covariate in all other analyses. For ANOVAs, univariate tests used Greenhouse-Geisser-corrected degrees of freedom^40^ (while corrected *df*s were used in analysis, uncorrected *df*s are presented in accordance with convention). Finally, following each grouping approach, convergence of approaches was also tested with a *χ*^*2*^ test on the Startle Group X Clinical Group contingency table.

## Results

### Sample Characterization

Ten participants (7.4%) were excluded from analyses due to missing startle data (poor data quality, technical failure, non-completion of the task), and another 13 participants (9.6%) could not be classified due to very low probability of responding to startle probes. The sample for all analyses below included 112 participants, who had a mean age of 32.4 (SD=10.1) and was predominantly female-identifying (N=84, 75%) and non-Hispanic white (N=91, 81.3%). In this sample, mean STAI-T score was 45.9 (SD= 12.2), mean BDI-II score was 13.7 (SD=10.9), mean PSWQ score was 25.7 (SD=8.7), and mean IIRS score was 44.7 (SD=21.5). Also in this sample as a whole, female sex at birth associated with decreased self-reported impairment (IIRS), *t*(103)=-2.1, *p*=.04, *d*=-0.5, and avoidance (BEAQ), *t*(103)=-2.5, *p*=.01, *d*=-.57, and also with increased worry (PSWQ-A), *t*=2.0, *p*=.046, *d*=.44.

### Manipulation Check: Task Ratings and Performance

To confirm that participants found task contexts evocative and were engaged, ratings and hit rate/ reaction times (RTs) were analyzed. For ratings, an effect of task context arose, *F*(4,444)=88.0, *p*<.001, 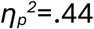, such that, in follow-up Bonferroni-corrected (n tests=10) paired-samples *t*-tests, participants rated avoidance (M=5.0, SD= 1.8), *t*(111)=5.0, *p*<.001, *d*=0.6, and escape (M=5.5, SD=1.6), *t*(111)=7.8, *p*<.001, *d*=0.9, as more unpleasant than neutral pictures (M=4.1, SD= 1.3) and not reliably different from each other, *t*(111)=2.7, *p*=.06, *d*=0.3; and, they also rated no-control cues (M=6.5, SD=1.5) as more unpleasant than escape cues, *t*(111)=5.6, *p*<.001, *d*=0.7, and disgust/violence images (M=7.1, SD=1.4) as more unpleasant than no-control cues, *t*(111)=3.3, *p*=.009, *d*=0.4. For performance, findings were similar to prior work such that participants 1) were generally and similarly successfully in executing avoidance (M=84.0%, SD=20.9) and escape (M=81.1%, SD=21.7) responses, *t*(111)= 1.5, *p*=.15, *d*=0.1, and 2) had similar RTs for avoidance (M=360.9, SD= 104.3) and escape (M=365.3, SD=105.8) responses, *t*(109)=0.5, *p*=.60, *d*=0.1.

### Blink Startle Reflex: General Task Effects

Results in the complete sample confirmed startle modulation across task contexts, *F*(2,222)=21.2, *p*<.001, 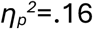, that replicated prior observations. Characterizing this modulation confirmed, first, that blink reactivity was potentiated in an no-control context as compared to an avoidance preparation context, *t*(110)=6.4, *p*<.001, *d*=0.8. Next, in this largely anxious treatment-seeking sample reactivity during escape preparation was also enhanced as compared to reactivity during avoidance preparation, *t*(110)=4.0, *p*<.001, *d*=0.5. Finally, reactivity was then further enhanced during no-control unpleasant anticipation as compared to escape preparation, *t*(110)=-2.5, *p*=.045, *d*=-0.3 (see **Figure 2**).

**Figure 2.**
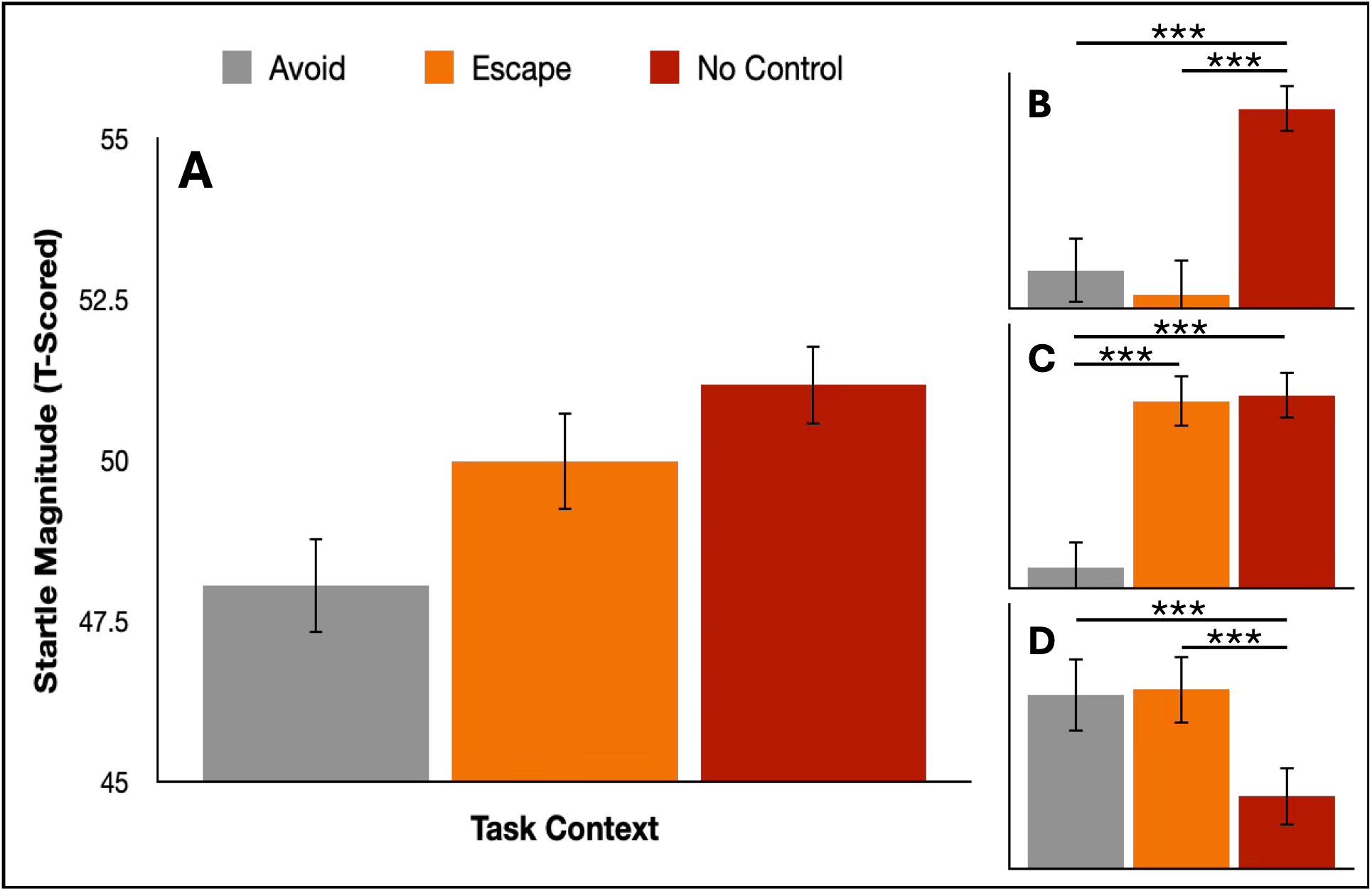
Startle modulation in the escape/ avoidance task. **Panel A** depicts mean blink amplitudes (T-scored within-subject relative to the person’s mean and standard deviation of all blinks) across avoid, escape, and no-control contexts in the whole sample. **Panel B, Panel C**, and **Panel D** depict modulation for action-sensitive, threat-sensitive, and context-insensitive groups, respectively. Error bars are 95% confidence intervals of the mean.

### Blink Startle Reflex: Classification and Relationship to Self-Report Clinical Variables

Classification of startle potentiation in no-control and escape contexts identified 35 individuals in an “action-sensitive” group, 45 in a “threat-sensitive” group, and 32 in a “context-insensitive” group (see **Figure 2**). Confirming context specificity of the difference between action- and threat-sensitive participants, while these groups differed by definition in startle reactivity during escape preparation, *t*(78)=7.5, *p*<.001, *d*=1.7, they did not differ in their reactivity in avoidance, *t*(78)=-1.0, *p*=.97, *d*=-0.2, or no-control, *t*(78)=-1.7, *p*=.29, *d*=-0.4, contexts. Next, context-insensitive subjects had reduced blink reactivity in a no-control context compared to action-sensitive, *t*(65)=-7.6, *p*<.001, *d*=-1.9, and threat-sensitive, *t*(75)=-6.4, *p*<.001, *d*=-1.5, participants, and they also had increased reactivity in an avoidance context as compared to action-, *t*(65)=5.1, *p*<.001, *d*=1.3, and threat-, *t*(75)= 6.4, *p*<.001, *d*=1.5, sensitive participants. In an escape context, context-insensitive participants had larger startle than an action-sensitive group, *t*(65)=6.5, *p*<.001, *d*=1.6, but did not differ from a threat-sensitive group, *t*(75)=-0.4, *p*>.99, *d*=-0.1. **Figure 2** depicts statistical patterns of modulation across contexts within each group.

After confirming profiles of reflex modulation, sample demographics and clinical differences were analyzed across groups (see **Table 1**). Regarding demographics, groups did not differ by age or race/ ethnicity but did differ in sex – such that there were fewer men in the context-insensitive group than in action-, *χ*^2^(1)= 6.0, *p*=.02, or threat-, *χ*^2^(1)=4.3, *p*=.04, sensitive groups. Given sex differences, analyses of symptom scores controlled for sex and revealed, first, that STAI-T scores were higher for threat-sensitive, *t*(78)=2.6, *p*=.03, *d*=0.6, and context-insensitive, *t*(65)=2.6, *p*=.03, *d*=0.7, groups than the action-sensitive group and they did not differ across context-insensitive and threat-sensitive participants, *t*(75)=0.2, *p*>.99, *d*=0.1. Additionally, PSWQ-A scores were higher in the threat-sensitive than the action-sensitive group, *t*(78)=2.5, *p*=.045, *d*=0.6, but they did not differ for context-insensitive participants compared to action-, *t*(65)=1.0, *p*=.94, *d*=0.3, or threat-, *t*(75)=-1.3, *p*=.60, *d*=-0.3, sensitive groups. Next, differences in BDI were driven by increased severity in the context-insensitive group compared to the action-sensitive group, *t*(65)=3.0, *p*=.01, *d*=0.7, while the threat-sensitive group did not differ from action-sensitive, *t*(78)=1.9, *p*=.17, *d*=0.4, or context-insensitive, *t*(75)=-0.3, *p*=.57, *d*=-0.3, groups. Finally, IIRS scores did not reliably differ for threat-sensitive and action-sensitive groups, *t*(78)=2.3, *p*=.08, *d*=0.5, or context-insensitive and threat-sensitive groups, *t*(75)=1.0, *p*>.99, *d*=0.2, but they did differ across context-insensitive and action-sensitive groups, *t*(65)=2.9, *p*=.01, *d*=0.7.

**Table 1.** Sample characteristics by startle group.

| **Demographics** | Action-Sensitive<br>N=35 | Threat-Sensitive<br>N=45 | Context-Insensitive<br>N=32 | $\chi^2$/ANOVA |
| --- | --- | --- | --- | --- |
| Sex (Female) | 23 (65.7%) | 32 (71.1%) | 29 (90.6%) | $\chi^2=6.1$, $p=.046^*$ |
| Race (White non-Hisp) | 25 (71.4%) | 38 (84.4%) | 28 (87.5%) | $\chi^2=3.3$, $p=.19$ |
| Age (M/(SD)) | 33.3 (11.2) | 32.2 (9.9) | 31.2 (9.4) | $F_{[2,109]}=0.2$, $p=.78$ |
| **Questionnaires** |  |  |  |  |
| STAI-T (M(SD)) | 40.9 (11.6)<sup>a</sup> | 48.0 (9.6)<sup>b</sup> | 48.3 (14.8)<sup>b</sup> | $F_{[2,108]}=4.5$, $p=.01^*$ |
| BDI (M(SD)) | 9.7 (8.8)<sup>a</sup> | 14.2 (8.9)<sup>a,b</sup> | 17.2 (14.1)<sup>b</sup> | $F_{[2,108]}=4.6$, $p=.01^*$ |
| PSWQ-A (M(SD)) | 22.9 (8.4)<sup>a</sup> | 27.8 (7.7)<sup>b</sup> | 25.9 (9.6)<sup>a,b</sup> | $F_{[2,108]}=3.1$, $p=.05^*$ |
| IIRS (M(SD)) | 37.3 (19.9)<sup>a</sup> | 47.2 (17.3)<sup>b</sup> | 49.4 (26.7)<sup>b</sup> | $F_{[2,108]}=4.7$, $p=.01^*$ |
| PCL (M(SD)) | 17.4 (17.9) | 22.7 (17.6) | 23.3 (22.4) | $F_{[2,108]}=1.2$, $p=.32$ |
| ASI (M(SD)) | 18.0 (11.1) | 21.0 (11.9) | 23.5 (17.1) | $F_{[2,108]}=1.5$, $p=.23$ |
| IUS-12 (M(SD)) | 31.4 (9.6)<sup>a</sup> | 35.1 (9.6)<sup>a,b</sup> | 37.6 (11.4)<sup>b</sup> | $F_{[2,108]}=3.1$, $p=.05^*$ |
| BEAQ (M(SD)) | 46.5 (13.5)<sup>a</sup> | 50.4 (9.6)<sup>a,b</sup> | 52.4 (17.0)<sup>b</sup> | $F_{[2,108]}=3.0$, $p=.05^*$ |

*Note:* Analyses of questionnaire scores are controlling for biological sex. $*p<0.05$

### Startle Differences across Groups based on Typical Clinical Classification

When participants were grouped by literature-based STAI-T and BDI-II cut-off scores as in prior work (rather than by startle modulation), n=38 individuals fell into a subthreshold symptoms group, n=58 into a clinically anxious (but not severely depressed) group, and n=16 into a severely depressed group. Given PSWQ differences identified above, exploring clinic-level worrying as an additional criterion further classified individuals with sub-threshold anxiety and depression into groups who did (n=12) or did not (n=26) have clinically elevated worry (worry consistently elevated in anxious and depressed groups). **Table 2** presents mean symptom levels across all questionnaires for each group.

**Table 2.** Clinical cutoff group characteristics.

| Group | Subthreshold<br>(n=26) | Worry Only<br>(n=12) | Clinical<br>Anxiety<br>(n=58) | Severe<br>Depression<br>(N=16) | $\chi^2$/ANOVA |
| --- | --- | --- | --- | --- | --- |
| Sex (Female) | 17 (65.4%) | 11 (91.7%) | 44 (75.9%) | 12 (75.0%) | $\chi^2=3.1$, $p=.38$ |
| Race (Non-hisp. white) | 21 (80.8%) | 9 (75.0%) | 46 (79.3%) | 15 (93.8%) | $\chi^2=2.1$, $p=.55$ |
| Age (M(SD)) | 34.2 (10.0) | 31.1 (11.2) | 32.9 (10.6) | 28.8 (7.4) | $F_{[3,108]}=1.1$, $p=.37$ |
| **Survey** |  |  |  |  |  |
| STAI-T<sub>(M(SD))</sub> | 31.1 (4.5)<sup>a</sup> | 34.8 (4.4)<sup>a</sup> | 49.6 (5.6)<sup>b</sup> | 64.6 (6.0)<sup>c</sup> | $F_{[3,108]}=162.1$<br>$\eta^2=.82^*$ |
| BDI<sub>(M(SD))</sub> | 3.8 (3.4)<sup>a</sup> | 6.4 (3.6)<sup>a</sup> | 13.9 (6.5)<sup>b</sup> | 34.3 (5.3)<sup>c</sup> | $F_{[3,108]}=110.0$,<br>$\eta^2=.75^*$ |
| PSWQ-<br>A<sub>(M(SD))</sub> | 14.6 (3.8)<sup>a</sup> | 26.6 (4.0)<sup>b</sup> | 28.3 (7.0)<sup>b</sup> | 33.8 (5.0)<sup>c</sup> | $F_{[3,108]}=45.6$,<br>$\eta^2=.56^*$ |
| IIRS<sub>(M(SD))</sub> | 26.8 (13.8)<sup>a</sup> | 29.8 (16.5)<sup>a</sup> | 47.5 (15.6)<sup>b</sup> | 75.1 (15.4)<sup>c</sup> | $F_{[3,108]}=37.5$,<br>$\eta^2=.51^*$ |
| PCL<sub>(M(SD))</sub> | 5.8 (7.1)<sup>a</sup> | 13.8 (13.4)<sup>a,b</sup> | 23.3 (17.4)<sup>b</sup> | 44.3 (17.9)<sup>c</sup> | $F_{[3,108]}=22.1$,<br>$\eta^2=.38^*$ |
| ASI<sub>(M(SD))</sub> | 10.1 (7.4)<sup>a</sup> | 19.1 (13.1)<sup>a,b</sup> | 21.1 (9.6)<sup>b</sup> | 38.6 (15.2)<sup>c</sup> | $F_{[3,108]}=24.3$,<br>$\eta^2=.40^*$ |
| IUS-12<sub>(M(SD))</sub> | 25.6 (7.4)<sup>a</sup> | 32.2 (8.9)<sup>a,b</sup> | 35.8 (8.4)<sup>b</sup> | 47.1 (7.9)<sup>c</sup> | $F_{[3,108]}=23.8$,<br>$\eta^2=.40^*$ |
| BEAQ<sub>(M(SD))</sub> | 41.3 (11.3)<sup>a</sup> | 41.9 (6.3)<sup>a,b</sup> | 50.5 (11.3)<sup>b</sup> | 65.8 (11.5)<sup>c</sup> | $F_{[3,108]}=18.3$,<br>$\eta^2=.34^*$ |

Note: Effect size ( $\eta^2$ ) rather than $p$ value shown since all $ps<.0001$ . $*p<.001$

After symptom-based classification, examining demographic characteristics revealed no differences in age, sex, or race/ ethnicity (see **Table 2**). For startle, a Clinical Group X Context interaction, *F*(6,216)=3.7, *p*=.002, 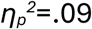, confirmed distinct modulation patterns as a function of different symptom profiles (**Figure 3**). Following this interaction, examining modulation within each group revealed that: a) a subthreshold group showed potentiation of blinks in a no-control context compared to avoid, *t*(25)=2.5, *p*=.049, *d*=0.7, and escape, *t*(25)=3.4, p=.005, *d*=0.9, contexts and no difference between avoid vs. escape contexts, *t*(25)=0.9, *p*>.99, 0.2; b) a worry-only group also exhibited startle potentiation in no-control, relative to both avoid, *t*(11)=3.4, *p*=.007, *d*=1.2, and escape, *t*(11)=3.4, *p*=.007, *d*=1.2, contexts and no difference between avoid and escape contexts, *t*(11)=0.1, *p*>.99, *d*=.01; c) a clinically anxious group showed potentiation in no-control, *t*(57)=6.0, *p*<.001, *d*=1.0, and escape, *t*(58)=5.6, *p*<.001, *d*=1.0, relative to avoid, contexts and no difference between no-control and escape contexts, *t*(58)=0.3, *p*>.99, *d*=0.1; and finally, d) a depressed group did not differ across no-control and avoid, *t*(15)=0.4, *p*>.99, *d*=0.1, escape and avoid, *t*(15)=-0.1, *p*>.99, *d*=-.04, or no-control and escape, *t*(15)=0.5, *p*>.99, *d*=0.2, contexts.

**Figure 3.**
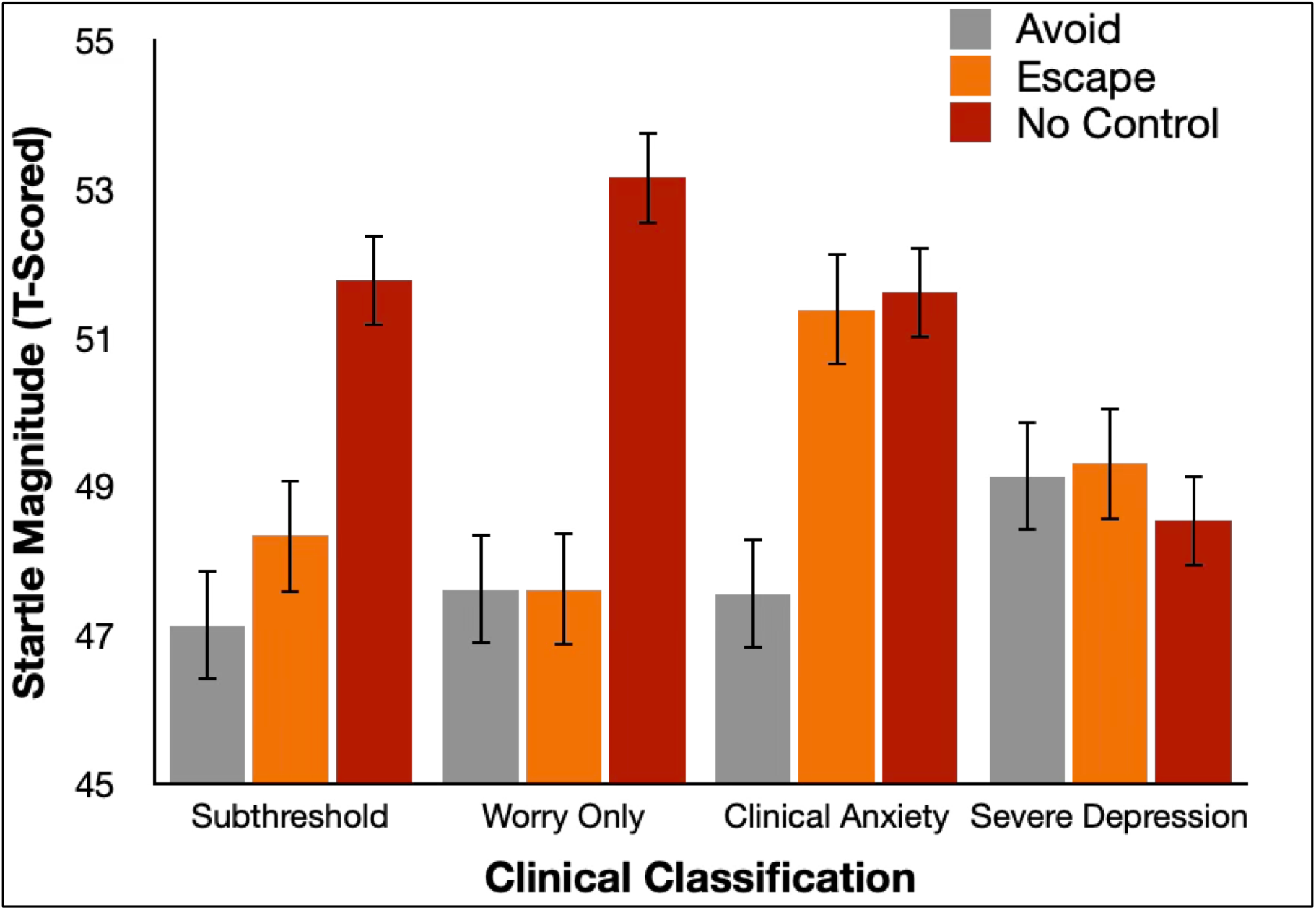
Startle modulation for subthreshold (PSWQ-A-, STAI-T-, BDI-), worry only (PSWQ-A+, STAI-T-, BDI-), clinical anxiety (STAI-T+, BDI-), and severe depression (BDI+) groups. Blinks are T-scored within-subject relative to that subject’s mean and standard deviation of all blinks. Error bars are 95% confidence intervals of the mean within each task condition.

Next, comparing groups within each task context revealed no group differences in an avoid context, *F*(3,108)=0.7, *p*=.58, 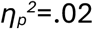, but differences in escape, *F*(3,108)=4.0, *p*=.01, 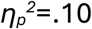, and no-control, *F*(3,101)=4.2, *p*=.008, 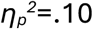, contexts. Differences were such that: a) in an escape context, an anxiety-only group had larger startle amplitudes than subthreshold, *t*(82)=2.8, *p*=.04, *d*=0.7, and (at a trend level) worry-only, *t*(68)=2.5, *p*=.07, *d*=0.8, groups, and; b) in a no-control context, severely depressed subjects had smaller blinks than did subthreshold, *t*(42)=-2.7, *p*=.045, *d*=-0.9, worry-only, *t*(26)=-3.2, *p*=.01, *d*=-1.2, and clinically anxious, *t*(72)=-2.9, *p*=.03, *d*=-0.8, groups.

### Convergence Tests

A Pearson chi-squared test revealed a significant association between startle profile-based and symptom cut-off-based groups, *χ*^*2*^(6)=27.3, *p*<.001. This was due to greater than 50% convergence: a) between an action-sensitive startle profile and both subthreshold and worry-only clinical groupings; b) between a threat-sensitive startle profile and an anxious clinical grouping, and; c) between a context-insensitive startle profile and a severely depressed clinical grouping. **Table 3** displays convergence of all groups.

**Table 3.** Convergence between clinical cutoff groups and startle profile groups.

|  | N | Action-sensitive | Threat-sensitive | Context-insensitive |
| --- | --- | --- | --- | --- |
| Subthreshold | 26 | 14 (53.9%)* | 5 (19.2%) | 7 (26.9%) |
| Worry Only | 12 | 7 (58.3%)* | 3 (25.0%) | 2 (16.7%) |
| Clinical Anxiety | 58 | 12 (20.7%) | 33 (56.9%)* | 13 (22.4%) |
| Severe Depression | 16 | 2 (12.5%) | 4 (25.0%) | 10 (62.5%)* |

\*Greater than 50% convergence

## Discussion

The goal of this work was to test if classifying individuals with varying levels of anxiety and depression based on their startle modulation in threat coping contexts captures clinical differences and/ or converges with conventional symptom-based profiling. Classifying by reflex potentiation in certain unpleasant exposure contexts revealed 1) two groups who differed specifically in their potentiation during escape preparation (inhibition during avoidance preparation and potentiation during uncontrollable anticipation were the same), and 2) a third group with disorganized modulation such that reflexes were apparently inhibited in an uncontrollable context but also less inhibited in an avoidance context. Building on our prior findings, comparing anxiety/ depression symptoms across startle-based profiles then revealed that: 1) a group with blink potentiation in uncontrollable *and* escape contexts (“threat-sensitive” group) had higher STAI-T anxiety and PSWQ-A worrying, but not BDI-II depression, than a group with potentiation in an uncontrollable context but *not* an escape context (“action-sensitive” group); and 2) a group with disorganized (“context-insensitive”) reflex modulation had higher BDI-II depression (but not worrying) than an action-sensitive group. Complementing reflex-based profiling, classifying individuals by symptoms then also showed that individuals low in STAI-T anxiety and BDI-II depression (but regardless of worrying) evinced action-sensitive-type modulation while individuals who were high in STAI anxiety but not severely depressed showed a threat-sensitive profile; and, those who *were* severely depressed had context-insensitive modulation. In concert with ∼54 – 63% agreement between startle- and symptom-based groupings, then, current findings extend evidence that an escape/ avoid preparation task captures clinically relevant change in underlying defense (“fight-flight-freeze”) system operation to guide treatment decisions.

Current analyses extend an approach laid out by Lang and colleagues^29^ to establishing psychiatric disorder biomarkers by testing neurobiological processing measures as clinical indicators rather than dependent variables – an approach which is entirely in line with initiatives like Research Domain Criteria (RDoC) that emphasize a need to guide treatment with objective indicators rather than subjective symptom reports.^7,8^ By extending this to defensive function in active threat contexts, current work is aligned with longstanding clinical doctrine that identifies avoidance-motivating defense system malfunctions as the core mechanism of anxiety, posttraumatic stress, and obsessive-compulsive disorders;^9,10^ and, with preclinical and clinical models that describe subtle dysregulations in defensive operation as more crucial than broad-based exaggerations.^16-19^ Inasmuch as an escape/ avoidance task is sensitive to context-specific defensive malfunction in many treatment seekers, it may be that this elicitation context can then one day be used to actually target this context-specific change – perhaps via pairing with precision neuromodulation^42^ – as an alternative to bluntly targeting putatively broad fear exaggeration as in frontline exposure-based therapies. In addition, converging evidence that an escape/ avoid task assay is also sensitive to a distinct pattern of defense dysregulation in depressed individuals converges with similar findings in other (script-driven emotional imagery^29,43-45^) tasks – and suggests that an indeed broad defensive shutdown (“defeat”^46,47^) in depressed individuals can also be detected by this measure. Importantly, this also implies that a different treatment might be needed for these individuals – one that, perhaps, seeks to *increase*, rather than regulate, defensive activation by targeting different brain areas during behavior challenge. Critically, increasing precision and individual tailoring of intervention in this way will be crucial to improving efficacy and also efficiency and tolerability of fear-targeting treatments^48,49^ – a critical goal inasmuch as current frontline approaches continue to be difficult for many treatment seekers and, thus, marked by significant non-response/ dropout rates.^50^

At the same time that current results extend the promise of our escape/ avoidance task, the 60% or less convergence of neurobiological and clinical profiles, variability of reactivity patterns within symptom profile groups, and variability in symptom severity within startle profile groups also confirm the need to keep improving reliability and predictiveness of this method. As one promising idea for improving this method, current and prior work have used normatively unpleasant images to standardize the elicitation context across subjects – but this also might undermine defensive evocativeness for participants who find these standard images less salient/ unpleasant. With this in mind, selecting images for personal/ diagnostic relevance in future work could lead to more reliable elicitation of defensive activation (and clinical aberrations therein) while also more closely paralleling exposure-based methods that build personalization into their protocols. As another candidate, future instantiations of our escape/ avoidance method might also seek ways to improve reliability of the startle measure (e.g., by conducting the task in acoustically optimized environments or adjusting loudness based on hearing tests) – an approach which could especially help reduce the startle non-response rate that is commonly observed in studies with this measure.^38^ Lastly, future work might also address unreliability on the clinic characterization side – such that, inasmuch as retrospective memory-based measures can be impacted by limited subjective insight into underlying processes or demand characteristics,^51-54^ future work might explore questionnaire batteries and/ or other sources of information as dependent variables rather than individual questionnaires.

In addition, some non-convergence may be due not to measure limitations, but the fact that blink startle is capturing unique, transdiagnostic variance that is important to psychiatric etiology but also not apparent in the subjective symptom domain. With this in mind, an additional future research need will be to determine if the escape/ avoidance task predicts other clinical variables perhaps more closely than does baseline self-report – with the most critical variables being treatment outcomes and tolerability patterns. To do this, future research should carry out longitudinal investigations that directly relate pre-treatment profiles of startle modulation in active threat coping contexts to post-treatment changes in anxiety, depression, and functional impairment. In addition, future work could test if directly modulating these profiles is more closely tied to functional changes than is self-report symptom-guided treatment – a result that would provide experimental manipulation-based confirmation that such profiles capture the underlying processing changes that drive impairment as causal mechanisms of anxiety and related disorders.

In addition to conceptual extensions described above, future research will need to address limitations of the present study. Most critically, future work must increase demographic – and especially racial/ ethnic – representativeness to test if current observations generalize beyond a predominantly white non-Hispanic female sample as represented here. Next, future research also must incorporate more thorough baseline clinical characterization to capture other clinical variables – e.g., disorder chronicity and treatment history – that were not analyzed here. Finally, future research should also continue to expand the comparison of treatment-seeking to non-treatment-seeking groups, and also the direct comparison of different diagnostic (e.g., primary posttraumatic stress vs. primary obsessive-compulsive disorder) groups, to confirm if current observations are truly dimensional/ transdiagnostic or, instead, are moderated by diagnostic/ treatment-seeking status.

The current study, then, provides support for continued investigation of blink startle profile in active threat coping contexts as a neurophysiological marker of defense system function that predicts clinical outcomes. Ultimately, clinical utility will be determined by the effectiveness of startle profiles to predict treatment response, which ultimately should be determined in controlled clinical trials. Such work should also include continued testing of how manipulating task features (e.g., personalized vs. standardized stimuli) impacts validity, and eaorts to reduce startle non-response rate for white noise bursts to increase utility for as many patients as possible. Studies on the neural correlates of blink startle responses^55,56^ may also help shape hypotheses on how startle profiles can inform clinical decision making. Ultimately, such research could lay groundwork for a precision medicine approach to treating clinical anxiety that increase effectiveness by increasing precision and directness in targeting underlying mechanisms rather than overlying symptoms.

## Footnotes

1 All analyses were also repeated in the sample of newly recruited individuals only, to evaluate degree to which results in this sample replicate our previous report.^13^Results are presented in supplemental materials

2 No participant had a No Control – Avoid score between 0 and 1, and no participant had an Escape – Avoid score between 0 and 1.

