## Supplemental Material for "Toward an Objective Assessment of Fight-or-Flight System Function in Anxiety Disorders: Startle Reflex Modulation in Threat Coping Contexts as a Candidate Neurophysiological Marker of Disrupted Defense System Operation"

To examine the consistency of patterns observed in the full combined-study sample and replicability of findings reported in our prior work (Sege et al., 2023), analyses were repeated with only newly recruited subjects included. All analytic procedures were identical to those described in the main article.

Additionally, analyses were repeated a third time with all treatment-seekers included (i.e., including individuals who were actively treatment-seeking, but not those who were not seeking treatment, from prior work). Descriptive patterns were very similar in this analysis as to those described next, and significances of effects were the same except where noted.

**Results**

*Sample Characterization*

Without participants from prior research included, the sample of 74 newly recruited treatment seekers with usable startle data had a mean age of 33.4 (SD=11.2) and was still mostly female-identifying (N=58, 78.4%) and non-Hispanic white (N=59, 79.7%). Mean STAI-T was 47.0 (SD= 12.5), mean BDI-II was 14.6 (SD=11.3), mean PSWQ-A was 26.8 (SD=8.3), and mean IIRS was 48.1 (SD=21.3). In this sample, female sex at birth was associated with decreased impairment (IIRS), $t(72)=-3.2$, $p=.002$, $d=-0.9$, and avoidance (BEAQ), $t(71)=-2.4$, $p=.02$, $d=-0.7$, but not with worry (PSWQ-A), $t(72)=0.8$, $p=.41$, $d=0.2$.

*Manipulation Check: Task Ratings and Performance*

Analysis of ratings still revealed an effect of task context, $F(4,292)=53.7$, $p<.001$, $\eta_p^2=.42$, such that, in follow-up Bonferroni-corrected (n tests=10) *t*-tests, participants rated avoid (M=4.9, SD= 1.8), $t(73)=3.3$, $p=.01$, $d=0.5$, and escape (M=5.4, SD=1.6), $t(73)=5.3$, $p<.001$, $d=0.8$, cues as more unpleasant than neutral pictures (M=4.1, SD= 1.4), and as not different from each other, $t(73)=2.1$, $p=.41$, $d=0.3$. They also still rated no-control cues (M=6.5, SD=1.5) as more unpleasant than escape cues, $t(73)=4.7$, $p<.001$, $d=0.7$, and disgust/violence images (M=7.1, SD=1.4) as marginally more unpleasant than no-control cues, $t(73)=2.8$, $p=.053$, $d=0.4$.[1] For performance, participants were still generally and similarly successfully in executing avoid (M=84.1%, SD=23.0) and escape (M=80.7%, SD=22.6) responses, $t(73)= 1.4$, $p=.17$, $d=0.2$, and they also still had similar RTs for avoid (M=365.6, SD= 117.0) and escape (M=359.7, SD=99.8) responses, $t(71)=0.5$, $p=.62$, $d=0.1$.

*Blink Startle Reflex: General Task Effects*

In the newly recruited sample, startle modulation across task contexts, $F(2,146)= 12.2$, $p<.001$, $\eta_p^2=.14$, was still in line with patterns in prior high-anxiety samples. First, blink

[1] In analysis with all treatment seekers, this comparison reached conventional significance, $t(95)=3.0$, $p=.03$, $d=0.4$

reflex reactivity in an uncontrollable unpleasant anticipation context (M=50.8; SD= 3.7) was potentiated relative to an avoidance preparation context (M=47.5; SD=4.7), $t(73)= 4.7$, $p<.001$, $d=0.8$. Second, for this actively treatment-seeking sample startle in the escape context (M=50.1; SD=4.6) was also enhanced relative to the avoidance context, $t(73)=3.7$, $p<.001$, $d=0.6$, and did not differ from the no-control context, $t(73)=-1.0$, $p=.95$, $d=-0.2$.

*Blink Startle Reflex: Classification and Relationship to Self-Report Clinical Variables*

In the sample of newly recruited participants, 21 individuals were classified into an "action-sensitive" startle modulation group, 30 into a "threat-sensitive" group, and 23 into a "context-insensitive" group. A context-specific difference between threat-sensitive and action-sensitive participants was maintained such that these groups differed in startle reactivity during escape preparation by definition, $t(49)=6.5$, $p<.001$, $d=1.9$, but did not differ in their reactivity in avoidance, $t(49)=-0.6$, $p>.99$, $d=-0.2$, or no-control, $t(49)=-1.8$, $p=.21$, $d=-0.5$, contexts. Next, context-insensitive subjects had reduced blink reactivity in a no-control context compared to action-sensitive, $t(42)=-6.6$, $p<.001$, $d=-2.0$, and threat-sensitive, $t(51)=-5.4$, $p<.001$, $d=-1.5$, participants, and they also had increased reactivity in an avoidance context as compared to action-, $t(42)=4.3$, $p<.001$, $d=1.3$, and threat-, $t(51)= 5.3$, $p<.001$, $d=1.5$, sensitive participants. In an escape context, context-insensitive participants had greater startle reactivity than an action-sensitive group, $t(42)=5.4$, $p<.001$, $d=1.6$, but they did not differ from a threat-sensitive group, $t(51)=-0.8$, $p>.99$, $d=-0.2$.

Modulation patterns within group were also like whole-sample patterns (see main article **Figure 2**). Thus: 1) in an action-sensitive group, startle was potentiated in no-control (M= 53.2; SD=3.3) relative to avoid (M=46.2; SD=4.5), $t(20)=7.6$, $p<.001$, $d=1.9$, and escape (M= 45.5; SD=3.0), $t(20)=8.3$, $p<.001$, $d=2.1$, contexts and did not differ across avoid vs. escape contexts, $t(20)=0.7$, $p<.001$, $d=0.2$; 2) in a threat-sensitive group, startle was potentiated in no-control (M=51.7; SD=2.8), $t(29)=7.9$, $p<.001$, $d=1.8$, and escape (M= 52.2; SD=3.6), $t(29)=8.5$, $p<.001$, $d=1.9$, compared to avoid (M=45.5; SD=3.8) contexts and did not differ in escape vs. no-control contexts, $t(29)=0.7$, $p>.99$, $d=0.2$, and; 3) in a context-insensitive group, startle was reduced in no-control (M=47.4; SD=2.5) compared to avoid (M=51.3; SD=3.5), $t(22)=-4.5$, $p<.001$, $d=1.1$, and escape (M=51.4; SD=4.1), $t(22)=4.7$, $p<.001$, $d=1.2$, contexts and did not differ across avoid and escape, $t(22)=0.2$, $p<.001$, $d=0.1$.

After confirming profiles of reflex modulation, sample demographics and clinical differences were analyzed across groups (see **Table S1**). As shown in **Table S1**, groups did not differ by age, race/ ethnicity, or biological sex. Next, Bonferroni-corrected symptom comparisons showed a non-significant difference in STAI-T score between threat-sensitive and action-sensitive groups, $t(49)=1.9$, $p=.21$, $d=0.6$, but a still-significant STAI-T difference between context-insensitive and action-sensitive groups, $t(42)=3.3$, $p=.005$, $d=1.3$ (context-insensitive and threat-sensitive participants still did not differ, $t[51]=1.9$, $p=.19$, $d=0.7$). Next, PSWQ-A scores did not differ across threat-sensitive and action-sensitive groups, $t(49)=2.1$, $p=.12$, $d=0.6$, context-insensitive and action-sensitive groups, $t(42)=1.7$, $p=.27$, $d=0.5$, or context-insensitive and threat-sensitive groups, $t(51)=-0.3$, $p>.99$, $d=-0.1$, groups. Regarding BDI, differences were still driven by increased severity in the context-insensitive group compared to the action-sensitive group, $t(42)=3.6$, $p=.002$, $d=1.1$, while the threat-sensitive group did not differ from action-sensitive, $t(49)=1.8$, $p=.25$, $d=0.5$, or context-insensitive, $t(51)=-2.1$, $p=.13$, $d=-0.6$, groups.[2] Finally, IIRS scores did not reliably differ for threat-sensitive and action-sensitive groups, $t(49)=1.3$, $p=.56$, $d=0.4$, or context-insensitive and threat-sensitive groups, $t(51)=2.1$, $p=.13$, $d=0.6$, but they did differ between context-insensitive and action-sensitive groups, $t(42)=3.2$, $p=.007$, $d=1.0$.[3]

*Startle Differences across Groups based on Typical Clinical Classification*

When participants were grouped by clinical cut-off scores as in main analyses, n=13 individuals fell into a subthreshold symptoms group, n=9 into a worry-only group, n=40 into a clinically anxious (but not severely depressed) group, and n=12 into a severely depressed group. Because of small group sizes and a lack of PSWQ-A differences in the new sample, in the following analyses subthreshold and worry-only groups were combined (n=22). **Table S2** presents mean symptom levels across all questionnaires for each group.

*Table S1. Sample characteristics by startle group.*

| **Demographics** | Action-Sensitive N=21 | Threat-Sensitive N=30 | Context-Insensitive N=23 | $\chi^2$/ANOVA |
|---|---|---|---|---|
| Sex (Female) | 16 (76.2%) | 22 (73.3%) | 20 (87.0%) | $\chi^2=1.5$, $p=.47$ |
| Race (White non-Hisp) | 15 (71.4%) | 24 (80.0%) | 20 (87.0%) | $\chi^2=1.6$, p=.44 |
| Age (M/(SD)) | 35.6 (13.2) | 34.2 (11.3) | 30.4 (8.6) | $F_{[2,71]}=1.3$, p=.28 |
| **Questionnaires** | | | | |
| STAI-T(M(SD)) | 40.7 (11.3)[a] | 47.5 (10.2)[a,b] | 52.0 (14.1)[b] | $F_{[2,70]}=5.3$,p=.007** |
| BDI(M(SD)) | 9.0 (7.3)[a] | 14.4 (9.0)[a,b] | 20.0 (14.5)[b] | $F_{[2,70]}=6.3$,p=.003** |
| PSWQ-A(M(SD)) | 23.5 (7.6) | 28.3 (8.2) | 28.0 (8.7) | $F_{[2,70]}=2.4$, p=.09^ |
| IIRS(M(SD)) | 39.9 (17.2)[a] | 47.7 (18.1)[a,b] | 56.0 (26.1)[b] | $F_{[2,70]}=5.1$,p=.009* |
| PCL(M(SD)) | 19.3 (18.2) | 21.5 (17.3) | 23.7 (22.0) | $F_{[2,70]}=0.4$, p=.70 |
| ASI(M(SD)) | 18.8 (10.0) | 20.3 (12.0) | 26.6 (17.2) | $F_{[2,70]}=2.1$,p=.13 |
| IUS-12(M(SD)) | 32.9 (9.0) | 37.3 (10.1) | 39.5 (11.2) | $F_{[2,70]}=2.5$, p=.09^ |
| BEAQ(M(SD)) | 47.0 (12.8) | 50.8 (10.9) | 54.8 (16.4) | $F_{[2,70]}=2.6$, p=.09^ |

*^p<.10; *p<0.05; **p<.01*

[2] In analysis with all treatment seekers, BDI scores were marginally higher in the context-insensitive group than in the threat-sensitive group, $t(95)=2.2$, $p=.096$, $d=0.5$

[3] Comparisons of clinical symptoms across startle groups continued to include sex as a covariate as in main analyses. Removing this covariate did not change statistical patterns in any instance

After symptom-based classification, examining demographic characteristics revealed no age, sex or race/ethnicity differences (see **Table S2**). For startle, a Symptom Group X Context interaction, $F(4,142)=4.3$, $p=.003$, $\eta_p^2=.11$, confirmed distinct modulation patterns in different symptom groups. Testing modulation in each group revealed that: a) a sub-threshold group had potentiated blinks in a no-control (M=52.2; SD=3.8) context compared to avoid (M=46.9; SD=5.5), $t(21)=3.4$, $p=.003$, $d=0.7$, and escape (M=48.1; SD=5.2), $t(21)=2.6$, p=.02, $d=0.6$, contexts and no difference between avoid vs. escape contexts, $t(21)=1.3$, $p=.22$, 0.3; b) a clinically anxious group showed potentiation in no-control (M= 50.9; SD=3.5), $t(39)=4.5$, $p<.001$, $d=0.7$, and escape (M=51.0; SD=4.2), $t(39)=4.6$, $p<.001$, $d=0.7$, relative to avoid (M=47.2; SD=3.6), contexts and no difference between no-control and escape contexts, $t(39)=-0.2$, $p=.87$, $d=.03$; and, c) a depressed group did not differ across no-control (M=48.0; SD=2.9) and avoid (M=49.3; SD=5.8), $t(11)=-0.6$, $p=.54$, $d=-0.2$, escape (M=50.5; SD=3.8) and avoid, $t(11)=0.6$, $p=.55$, $d=0.2$, or (at a conventional reliability level) no-control and escape, $t(11)=-1.9$, $p=.08$, $d=0.6$, contexts.

Next, comparing groups within each task context revealed no group differences in an avoid context, $F(2,71)=1.2$, $p=.32$, $\eta_p^2=.03$, but groups did differ in startle reactivity in escape, $F(2,71)=3.1$, $p=.05$, $\eta_p^2=.08$, and no-control, $F(2,71)=5.5$, $p=.006$, $\eta_p^2=.13$, contexts. Differences were such that: a) in an escape context, an anxiety-only group had larger startle amplitudes than a subthreshold group, $t(60)=2.4$, $p=.05$, $d=0.6$, and; b) in a no-control context, severely depressed subjects had smaller blinks than did subthreshold, $t(32)=-3.3$, $p=.005$, $d=-1.2$, and clinically anxious, $t(50)=-2.5$, $p=.047$, $d=-0.8$, groups.

*Table S2. Clinical cutoff group characteristics*

| **Group** | Subthreshold (n=22) | Clinical Anxiety (n=40) | Severe Depression (N=12) | $\chi^2$/ANOVA |
|---|---|---|---|---|
| Sex (Female) | 18 (81.8%) | 31 (77.5%) | 9 (75.0%) | $\chi^2=0.3$, p=.88 |
| Race (Non-hisp. white) | 18 (81.8%) | 30 (75.0%) | 11 (91.7%) | $\chi^2=1.7$, p=.43 |
| Age (M/(SD)) | 34.7 (11.7) | 34.2 (11.5) | 28.5 (7.8) | $F_{[2,71]}=1.4$, p=.25 |
| **Survey** | | | | |
| STAI-T (M(SD)) | 31.9 (3.5)[a] | 49.5 (5.6)[b] | 66.3 (3.9)[c] | $F_{[2,71]}=210.3$, $\eta^2=.86$* |
| BDI (M(SD)) | 4.8 (3.6)[a] | 14.0 (6.5)[b] | 34.8 (6.1)[c] | $F_{[2,71]}=107.5$, $\eta^2=.75$* |
| PSWQ (M(SD)) | 19.9 (6.7)[a] | 28.7 (7.3)[b] | 33.6 (5.5)[c] | $F_{[2,71]}=18.6$, $\eta^2=.34$* |
| IIRS (M(SD)) | 33.1 (16.1)[a] | 48.0 (16.2)[b] | 75.7 (17.7)[c] | $F_{[2,71]}=26.1$, $\eta^2=.42$* |
| PCL (M(SD)) | 11.2 (11.9)[a] | 20.9 (17.1)[b] | 42.8 (19.0)[c] | $F_{[2,71]}=15.0$, $\eta^2=.30$* |
| ASI (M(SD)) | 14.3 (10.6)[a] | 20.8 (9.0)[b] | 39.2 (16.8)[c] | $F_{[2,71]}=20.1$, $\eta^2=.36$* |
| IUS-12 (M(SD)) | 29.5 (8.7)[a] | 37.2 (8.7)[b] | 48.8 (6.0)[c] | $F_{[2,71]}=20.9$, $\eta^2=.37$* |
| BEAQ (M(SD)) | 42.8 (9.9)[a] | 50.9 (12.4)[b] | 65.3 (10.4)[c] | $F_{[2,71]}=14.6$, $\eta^2=.30$* |

*Note:* Effect size ($\eta^2$) rather than *p* value shown since all *p*s<.0001. *$p<.001$

*Convergence Tests*

A Pearson chi-squared test revealed a significant association between startle profile-based and symptom cut-off-based groups, $\chi^2(4)=25.3$, $p<.001$. This was due to greater than 50% convergence: a) between an action-sensitive startle profile and a subthreshold symptom grouping; b) between a threat-sensitive startle profile and an anxious clinical grouping, and; c) between a context-insensitive startle profile and a severely depressed clinical grouping. **Table S3** displays the convergence of all groups.

*Table S3. Convergence between clinical cutoff groups and startle profile groups.*

| | N | Action-sensitive | Threat-sensitive | Context-insensitive |
|---|---|---|---|---|
| Subthreshold | 22 | 13 (59.1%)* | 5 (22.7%) | 4 (18.2%) |
| Clinical Anxiety | 40 | 8 (20.0%) | 22 (55.0%)* | 10 (25.0%) |
| Severe Depression | 12 | 0 (0.0%) | 3 (25.0%) | 9 (75.0%)* |

**Greater than 50% convergence*

**Summary**

While sub-sample tests largely replicate main analyses, in tests of symptom differences across startle response groups the comparison of trait anxiety between action-sensitive and treat-sensitive participants was in the same direction but no longer reliably significant. Nonetheless, effect size for this effect was the same as in main analyses, suggesting a medium-sized effect that was no longer detected due to reduced power – and, potentially, altered distributional characteristics due to less representation of individuals truly low in trait anxiety. In all, this result again supports the need for continued research into how to improve reliability and sensitivity of the escape/ avoidance task (e.g., via personalization of task stimuli) as well as testing if effect sizes for group comparisons of other clinical variables (e.g., treatment outcome) are larger than for baseline symptom levels.

Beyond the distinction described above, all other results replicated those from the main analyses – including similar patterns of convergence between startle-based and symptom-based profiles – and thus further support conclusions in the main article. Importantly, examining startle modulation patterns across symptom-defined groups also entirely replicated patterns from our prior work (Sege et al., 2023) – such that a subsequently recruited sample of participants who were treatment-seeking but also low in trait anxiety had an action-sensitive-type pattern of startle modulation as a group, treatment seekers who were high in trait anxiety evinced threat-sensitive-type modulation as a group, and severely depressed treatment seekers had overall reduced modulation of startle. This result replicates the promise of the escape/ avoidance task as a tool for detecting clinically meaningful change in defensive system operation – and, again, supports continued research to develop this task for ultimate deployment as a clinical assessment tool.
